# Dynamics of an incoherent feedforward loop drive ERK-dependent pattern formation in the early *Drosophila* embryo

**DOI:** 10.1101/2023.03.09.531972

**Authors:** Emily K. Ho, Harrison R. Oatman, Sarah E. McFann, Liu Yang, Heath E. Johnson, Stanislav Y. Shvartsman, Jared E. Toettcher

## Abstract

Positional information in developing tissues often takes the form of stripes of gene expression that mark the boundaries of a particular cell type or morphogenetic process. How stripes form is still in many cases poorly understood. Here we use optogenetics and live-cell biosensors to investigate one such pattern: the posterior stripe of *brachyenteron (byn)* expression in the early *Drosophila* embryo. This *byn* stripe depends on interpretation of an upstream signal – a gradient of ERK kinase activity – and the expression of two target genes *tailless (tll)* and *huckebein (hkb)* that exert antagonistic control over *byn*. We find that high or low doses of ERK signaling produce either transient or sustained *byn* expression, respectively. These ERK stimuli also regulate *tll* and *hkb* expression with distinct dynamics: *tll* transcription is rapidly induced under both low and high stimuli, whereas *hkb* transcription converts graded ERK inputs into an output switch with a variable time delay. Antagonistic regulatory paths acting on different timescales are hallmarks of an incoherent feedforward loop architecture, which is sufficient to explain transient or sustained *byn* dynamics and adds temporal complexity to the steady-state model of *byn* stripe formation. We further show that an all-or-none stimulus can be ‘blurred’ through intracellular diffusion to non-locally produce a stripe of *byn* gene expression. Overall, our study provides a blueprint for using optogenetic inputs to dissect developmental signal interpretation in space and time.

## Introduction

Cells in a developing embryo must obtain information about their location to enact appropriate morphogenetic and fate decisions. In many developmental contexts this positional information takes the form of stripes of gene expression that subdivide a coordinate axis into distinct regions. Stripe formation requires precise spatial coordination of activating and inhibiting inputs to delineate boundaries. This complex signal processing has spurred decades of study of both natural and engineered stripe-forming biological systems (Schaerli et al. 2014; Stanojevic et al. 1991; Tabor et al. 2009). However, in many cases, how a stripe is produced by simple asymmetric inputs is still incompletely understood.

One of the simplest and best-studied developmental stripes is that of *brachyenteron (byn)* expression in the posterior region of the pre-gastrulation *Drosophila* embryo. *Byn* is an essential gut regulator and homolog of vertebrate *Brachyury* (Kispert et al. 1994; Singer et al. 1996; Kusch and Reuter 1999; McFann et al. 2022). It is primarily controlled by just a single input: activation of the Torso receptor tyrosine kinase at the embryo’s poles which produces a gradient of active, doubly-phosphorylated ERK (dpERK), known as the terminal pattern (Casanova and Struhl 1989; Lu et al. 1993; Sprenger and Nüsslein-Volhard 1992). ERK then inactivates the transcriptional repressor Capicua (Cic), leading to the expression of two essential transcription factors, *tailless* (*tll*) and *huckebein* (*hkb*) (**Figure 1A**) (Coppey et al. 2008; Jiménez et al. 2000; Strecker et al. 1986; Weigel et al. 1990; Pignoni et al. 1990; Bronner and Jackle 1991; Keenan et al. 2020; Patel et al. 2021). *Tll* is sensitive even to low doses of ERK and thus is expressed broadly, whereas *hkb* requires a high dose and thus is expressed at only the most posterior positions (Greenwood and Struhl 1997; Ghiglione et al. 1999). Expression of *byn* is positively regulated by Tll and repressed by Hkb, and so *byn* forms a stripe at low doses of ERK signal (Kispert et al. 1994).

**Figure 1:**
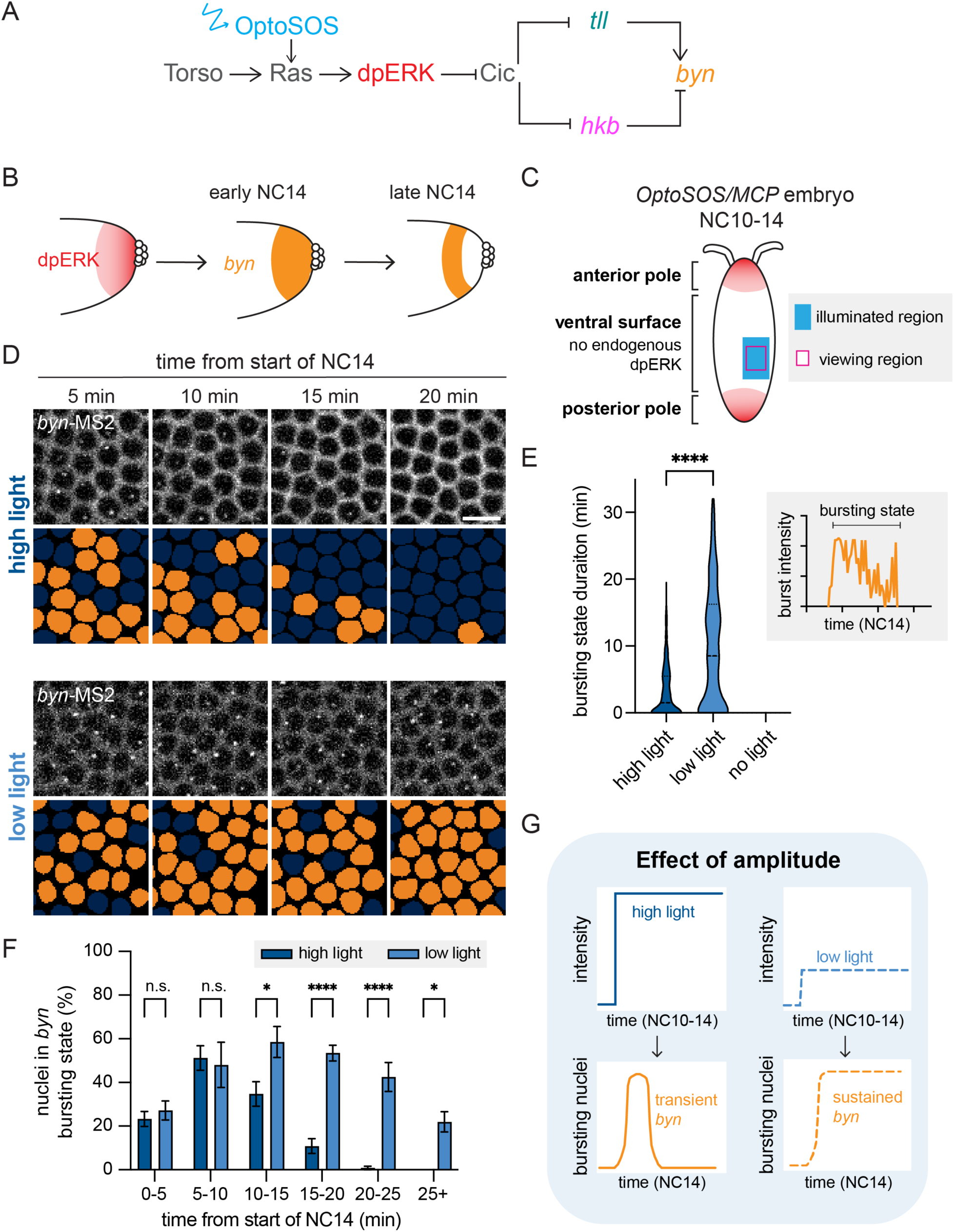
High and low amplitude optoSOS inputs yield transient and sustained *byn* transcription. **(A)** Diagram of the genetic circuit regulating *byn* expression. OptoSOS (blue arrow) activates the circuit at the level of Ras. **(B)** *byn* dynamics in NC14 relative to the endogenous dpERK gradient. **(C)** Schematic of blue light stimulation on the ventral side of wild-type OptoSOS/MCP embryos. The blue light is applied starting in NC10 to a region on one half of the ventral surface, away from the endogenous ERK activity at the poles. In NC14, *byn* transcription is imaged in a viewing window within the illuminated region. **(D)** *byn*-MS2 bursting during NC14 upon high or low blue light stimulation. In addition to the MS2 bursts, the membrane localized OptoSOS is visible in the same channel. Orange nuclear masks show nuclei with a burst in corresponding frames. Scale bar is 10 µm. **(E)** Time spent in *byn* bursting state under high, low, and no light. Inset shows that the bursting state is defined from the first to the last detectable MCP/MS2 focus. The bursting state is significantly longer under low light by ANOVA and Tukey’s post hoc test. n > 190 nuclei per condition **(F)** Proportion of nuclei in the *byn* bursting state through NC14. Under low light, significantly more nuclei remain in the bursting state after the first 10 minutes of NC14. Two-way ANOVA and Sidak’s post hoc test. Mean ± SEM, n = 4 embryos in each condition. **(G)** Schematic showing the effect of blue light intensity on *byn* expression dynamics.

Although the basic principles underlying *byn* stripe formation are well understood, some experimental results cannot be explained by the canonical model of steady-state regulation by *tll* and *hkb*. First, the pattern of *byn* expression is dynamic: it is initially expressed as a cap covering the entire posterior domain, only resolving into a stripe in late nuclear cycle 14 (NC14) (**Figure 1B**) (Kispert et al. 1994). Recently, Keenan et al. (2022) showed that these dynamics derive from changes in *byn* transcription: there is a domain of transient *byn* transcription at the pole and a domain of sustained *byn* transcription at interior positions. The source of these *byn* dynamics is not known, nor is it clear how *byn* might be expressed in the same posterior domain as its negative regulator *hkb*. Second, recent optogenetic studies demonstrated that replacing the terminal gradient by a strong all-or-none light input was sufficient to fully rescue embryogenesis (Johnson et al. 2020). This all-or-none input was observed to eliminate spatial differences in *tll* and *hkb* expression, such that the two genes were expressed in fully overlapping domains. How can a uniform input correctly pattern *byn*?

The architecture of the *byn* circuit provides a clue as to what may be missing from our understanding of *byn* regulation. The *byn* circuit has the general structure of an incoherent feedforward loop: a single input (dpERK) takes two paths (*tll* and *hkb*) with opposite signs to regulate a single output (**Figure 1A**) (Alon 2007). Incoherent feedforward loops can generate both steady-state stripes (Schaerli et al. 2014) and dynamic pulses (Mangan and Alon 2003), but the differences in the delay time through each path is a crucial parameter for the incoherent feedforward loop’s function. Whether the *tll* and *hkb* branches might exhibit different delays in the *byn* circuit is unknown and poses a conceptual challenge, given that the same ERK-dependent transcriptional regulator (Cic) mediates signal transfer to both target genes.

Here, we set out to examine dynamic transmission through the gene network regulating *byn* expression by combining optogenetic stimulation with endogenously-tagged transcriptional reporters at multiple network nodes. Optogenetics has proven to be a powerful approach to interrogate developmental circuits; the precision of light patterning enables the experimentalist to systematically vary the spatial and temporal parameters of an input while monitoring outputs ranging from gene expression to morphogenesis (McFann et al. 2021; Singh et al. 2022; Huang et al. 2017; Guglielmi et al. 2015). In particular, measuring responses after acute changes in stimulus can provide information about the timescale with which signals are transmitted through a developmental gene network (Singh et al. 2022; Patel et al. 2021; Keenan et al. 2020). We find that low- and high-amplitude optogenetic ERK stimuli are dynamically interpreted to generate either transient or sustained *byn* transcription. These dynamics are also reflected in the relative timing of *tll* and *hkb* transcription, with *tll* transcription occurring rapidly (< 5 min) but *hkb* transcription exhibiting a dose-sensitive delay of up to 1 hr post-illumination. Our data are consistent with a dose-dependent shift from transient to sustained *byn* expression dynamics that depends on the relative timing of Tll and Hkb protein accumulation. Finally, we show that a uniform input can also produce a *byn* stripe in space at a site near, but not overlapping with, the input stimulus, revealing a role for protein diffusion and mixing in the spatial interpretation of the terminal pattern.

## Results

### High and low optogenetic ERK inputs alter the dynamics of *brachyenteron* expression

Prior studies have shown that posterior *byn* expression is dynamic and forms two different domains (Kispert et al. 1994; Keenan et al. 2022). At the posterior pole, *byn* expression occurs in a transient pulse at the beginning of nuclear cycle 14 (NC14), whereas at more interior positions, *byn* is sustained longer into NC14 (**Figure 1B**). We first sought to test whether optogenetic inputs to ERK signaling are sufficient to reproduce these *byn* dynamics without crosstalk from the endogenous terminal pattern. We generated *Drosophila* embryos that expressed the OptoSOS system for Ras/ERK pathway activation (Johnson et al. 2017; Toettcher et al. 2013) and a fluorescent MCP protein for visualizing transcription of MS2-tagged transcripts (Garcia et al. 2013). These OptoSOS/MCP embryos also contained MS2 stem loops in the endogenous *byn* 5’UTR (*byn*-MS2) (Keenan et al. 2022). To avoid overlap with the endogenous terminal system, we stimulated a region on the embryo’s ventral surface that is sufficiently far from the pole to lack *byn*, *tll*, or *hkb* transcription when embryos are grown in the dark, but where *tll*, *hkb*, and *byn* expression could be readily induced by optogenetic stimulation (**Figure S1**). The light input was limited to the right half of the embryo (**Figure 1C**), providing an internally-controlled, unilluminated region at the same anterior-posterior and dorsoventral position. Blue light was delivered beginning in nuclear cycle 10 (NC10), coinciding with the start of endogenous terminal signaling, and continuing through NC14, a duration of roughly 90 min (Coppey et al. 2008).

We varied the intensity of the blue light dose to identify intensities that would produce *byn* responses similar to those endogenous responses observed by Keenan et al. (2022). Under a high-intensity blue light dose (“high light”), *byn* was expressed in a transient pulse, with MCP-labeled transcriptional foci detected for only 10-15 min after the start of NC14 (**Figure 1D**, top, **Movie S1**). This input was a similar intensity as that used previously to rescue the terminal pattern (Johnson et al. 2020). In contrast, an 8-fold lower low light dose (“low light”) produced *byn* transcription that was maintained in NC14 for 25 min or more (**Figure 1D**, bottom, **Movie S1**). To quantify these responses, we measured the number of bursting nuclei from a region of fixed size (**Figure S2A-B**), and we defined a nucleus as being in a “bursting state” for all time points between the first and last appearance of an MCP focus within a nuclear cycle (**Figure 1E**, inset). Quantification from multiple embryos confirmed that low-intensity light stimulation drove *byn* bursts over a significantly longer duration than high light (**Figure 1E**) and that the bursting state under low light persisted significantly later into NC14 (**Figure 1F**). Together, these data confirm that two different levels of light-induced ERK activation are filtered into distinct *byn* dynamics, producing transient and sustained transcriptional responses that closely resemble *byn*’s endogenous dynamics at the posterior pole (**Figure 1G**).

### *brachyenteron* dynamics are reflected in the kinetics of *tailless* and *huckebein*

We hypothesized that the differences in *byn* transcription in each light condition would be reflected in the expression of *tll* and *hkb*, *byn*’s well-characterized positive and negative regulators (**Figure 2A**). We applied the same high and low light stimuli to OptoSOS/MCP embryos whose endogenous *tll* or *hkb* genes contained MS2 stem loops in the 5’ UTR (*tll*-MS2 and *hkb*-MS2) (Keenan et al. 2022). These endogenously-tagged MS2 reporters differ from the enhancer sequences used in prior studies (Johnson et al. 2020; Keenan et al. 2020) and are expected to better reflect the full regulatory environment of each gene. We then analyzed transcription of each ERK-responsive gene over time (**Figure S2A-B**). Both high and low stimuli induced *tll* transcription in NC14, consistent with the standard model whereby *tll* transcription is triggered even by weak ERK stimuli (**Figure 2B**) (Greenwood and Struhl 1997). We also found that both high and low light could induce *hkb* transcription, with four out of seven embryos under low light exhibiting *hkb* expression in NC14 (**Figure 2B**, **Figure S1A**). This result was unexpected because *byn* is sustained in NC14 under low light conditions (**Figure 1D**) and *hkb* is a known repressor of *byn* expression. Moreover, *hkb* is thought to only be induced by strong ERK stimuli. The transcription of both *tll* and *hkb* declined after the first 15 minutes of NC14 (**Figure S2C-D**), consistent with a loss of input sensitivity of both genes that is also observed for their endogenous expression patterns at the posterior pole (Keenan et al. 2022).

**Figure 2:**
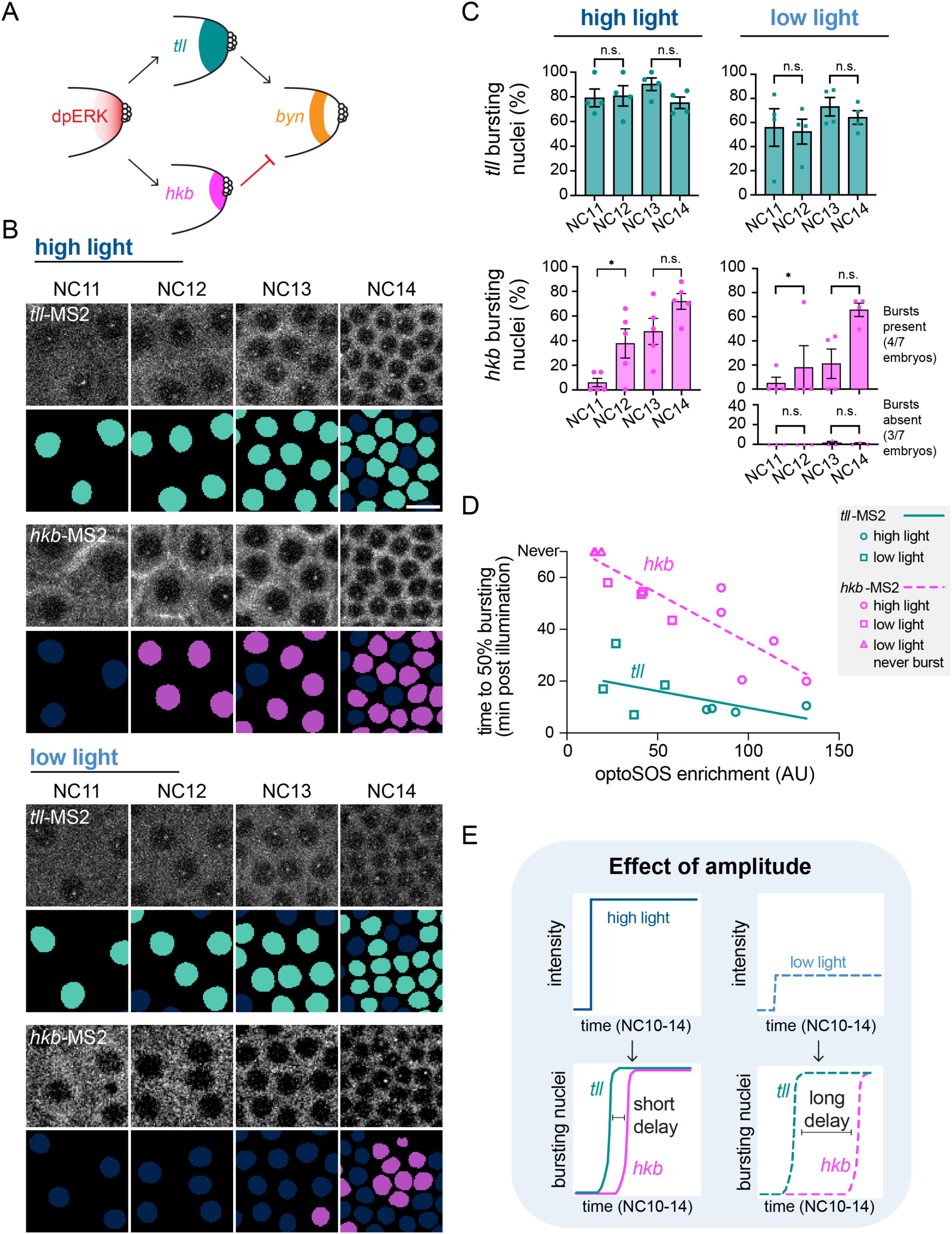
The onset of *tll* transcription in response to OptoSOS is rapid whereas the onset of *hkb* transcription is delayed and dose-dependent. **(A)** In the endogenous posterior pattern, *byn* is expressed in the low-ERK, *tll*-only region. **(B)** *tll-* and *hkb*-MS2 bursting during NC11-14 upon high or low blue light stimulation. Nuclear masks (*tll* in teal, *hkb* in magenta) show nuclei with a burst in corresponding frames. Scale bar is 10 µm. **(C)** Proportion of nuclei in the bursting state in each nuclear cycle for *tll* and *hkb* under high and low light. Under low light, *hkb* showed two different responses which are plotted on separate graphs. Each point represents one embryo and the mean ± SEM for each condition is shown. n = 4-7 embryos for each condition. Significance from ANOVA with Sidak’s post hoc test. **(D)** Plot relating optoSOS membrane enrichment as a proxy for ERK activity level to time to 50% bursting, measured from the start of illumination. Each point represents one embryo (*tll* in teal, *hkb* in magenta). The different shaped points show that embryos from the same illumination condition reach 50% bursting at similar but not identical times, depending on OptoSOS level. The line shows the linear regression of all points. **(E)** Schematic showing the effect of signal amplitude on *tll* and *hkb* dynamics. The delay between *tll* and *hkb* expression onset increases with decreasing signal amplitude.

Although high and low light stimuli drove similar *tll* and *hkb* transcription in NC14, the expression of these two genes were quite different in earlier nuclear cycles (NC11-13) (**Movie S2**). For *tll*, MS2 transcriptional foci appeared rapidly in NC11, and most nuclei maintained bursting states during NC11-14 (**Figure 2B-C**, teal). In contrast, expression of *hkb* was more delayed. Under high light, *hkb*-MS2 transcriptional foci were largely absent in NC11, but the proportion of nuclei in a bursting state significantly increased in NC12 and remained high in NC13 and 14 (**Figure 2B-C**, magenta). The delay between illumination and *hkb* transcription was much longer under low light, where *hkb* transcription was low or absent until NC14 (**Figure 2B-C**, magenta). These results reveal that *tll* and *hkb* exhibit different temporal expression dynamics in response to ERK activation, with *hkb* transcription delayed relative to *tll*. They also identify *hkb* as a factor whose transcriptional dynamics change as a function of ERK input strength. These results also differ from our prior observations using enhancer-based MS2 reporters (Johnson et al. 2020; Keenan et al. 2020), where we generally observed much weaker transcription in fewer nuclei and were unable to detect a difference in timing between *tll* and *hkb*. These differences may point to the importance of performing measurements at endogenous transcriptional loci, particularly when assessing gene expression dynamics.

We next set out to relate ERK inputs to the transcriptional delay in *tll* and hkb gene expression for individual embryos. OptoSOS expression can vary between embryos due to variability in Gal4/UAS-based control, so even a single light dose may produce different degrees of pathway stimulation. To account for this variability, we reasoned that embryo-specific inputs can be quantified by measuring the light-induced increase in SSPB-tagRFP-SOS^cat^ membrane fluorescence. By measuring the timepoint at which at least 50% of nuclei entered the bursting state as a function of each embryo’s SSPB-tagRFP-SOS^cat^ membrane recruitment, we were thus able to quantify the delays in *tll* and *hkb* expression over a continuum of ERK activity levels (**Figure 2D**). We found that the onset of *tll* transcription was rapid in nearly all embryos, reaching 50% bursting within as few as 8 minutes after the start of illumination, and exhibited weak scaling with the level of OptoSOS recruitment (**Figure 2D**, teal). In contrast, the onset of *hkb* expression occurred later than *tll* at all signal amplitudes and scaled strongly with ERK signal amplitude, ranging from 20 min to 60 min depending on membrane OptoSOS levels (**Figure 2D**, magenta). In the 3 embryos with the lowest ERK activity, *hkb* was never expressed. Taken together, these data demonstrate that nuclei of early *Drosophila* embryos convert the amplitude of ERK signaling into differences in the relative timing of *tll* and *hkb* transcription.

How might the relative delay in *tll* and *hkb* transcription affect their appearance at the protein level? To address this question we endogenously-tagged *tll* and *hkb* with the LlamaTag system to rapidly visualize protein-level accumulation of these transcription factors (Bothma et al. 2018) (**Figure S3A**). We applied our high light stimulus from NC10 onward (**Figure S3B**), a condition under which *tll* expression precedes *hkb* by ~10 min and a pulse of *byn* expression is observed in early NC14. We found that while nuclear Tll protein was detected both 10 min (early NC14) and 25 min into NC14 (late NC14), nuclear Hkb protein was only detected in late NC14 (**Figure S3B-D**). These data confirm that the relative delay between *tll* and *hkb* transcription is also reflected at the protein level for both factors. Further, they show that there is a large increase in Hkb protein in mid-NC14 that occurs around the same time as repression of *byn* begins.

Together, these results suggest a conceptual model for how ERK signals of different amplitudes produce either transient or sustained dynamics of *byn* expression (**Figure 2E**). When ERK activity is high, *tll* expression precedes *hkb* expression by a short but important window, leading to the pulse of Tll-mediated *byn* expression in early NC14 before the accumulation of Hkb repressor protein. When ERK is lower, *tll* expression is still rapidly initiated but *hkb* expression is substantially delayed (or, at sufficiently low ERK doses, altogether absent). The resulting Hkb protein levels do not rise until too late in NC14, leading to sustained *byn* expression. At the core of this model is the observation that differences in ERK dose are converted into temporal differences in *hkb* transcription onset.

### Time varying optogenetic inputs further validate the model of *brachyenteron* regulation

Our conceptual model holds that *tll* and *hkb* are induced by ERK after different time delays, producing distinct *byn* dynamics. This model has so far been tested only under a simple stimulus regime: a step-change in light delivered at NC10 and maintained until gastrulation. A strength of the optogenetic approach is that stimuli can be varied over time, either by switching between light and dark conditions or by varying the developmental time at which the stimulus is applied. Importantly, such dynamic experiments are possible because transcriptional repression by Capicua (Cic) is restored within minutes of turning off an optogenetic input to the ERK pathway (Patel et al. 2021).

We first applied a pulse of light to interrogate the consequences of the delay between *tll* and *hkb* expression. If a high light stimulus is delivered as a pulse that is sufficiently long to express *tll* but removed before *hkb* expression is initiated, it should be possible to express the Tll activator without Hkb-mediated repression, thereby shifting *byn* expression from transient to sustained. Importantly, this prediction differs from the canonical model, where a high ERK input would give rise to both *tll* and *hkb* expression, leading to *byn* repression. We applied high light to OptoSOS/MCP embryos as a pulse from NC10-13, a duration of roughly 45 min (**Figure 3A**, left), and compared *byn*-MS2 expression to our previous experiment where we applied a continuous stimulus beginning in NC10 (**Figure 1**). We found that these differences in stimulus dynamics had a profound effect on *byn* expression (**Figure 3A**, right). In pulse-stimulated embryos, *byn* expression in NC14 was sustained, persisting for over 25 minutes in nearly all illuminated nuclei (**Figure 3A-B**, **Movie S3**). In contrast, continuous high light led to a brief pulse of *byn* expression during the first 15 min of NC14 in fewer than half of the illuminated nuclei (**Figure 3A**, top). In summary, two ERK-activating stimuli of the same amplitude but different durations produce distinct dynamics of *byn* expression, consistent with an important role for delayed *hkb* expression in repressing *byn*. This result is also consistent with our prior observations concerning the role of Hkb in repressing *snail* during ventral furrow formation (McFann et al. 2021), where we found that sustained illumination over a time period that includes late NC13/early NC14 was most effective in suppressing ventral furrow invagination.

**Figure 3:**
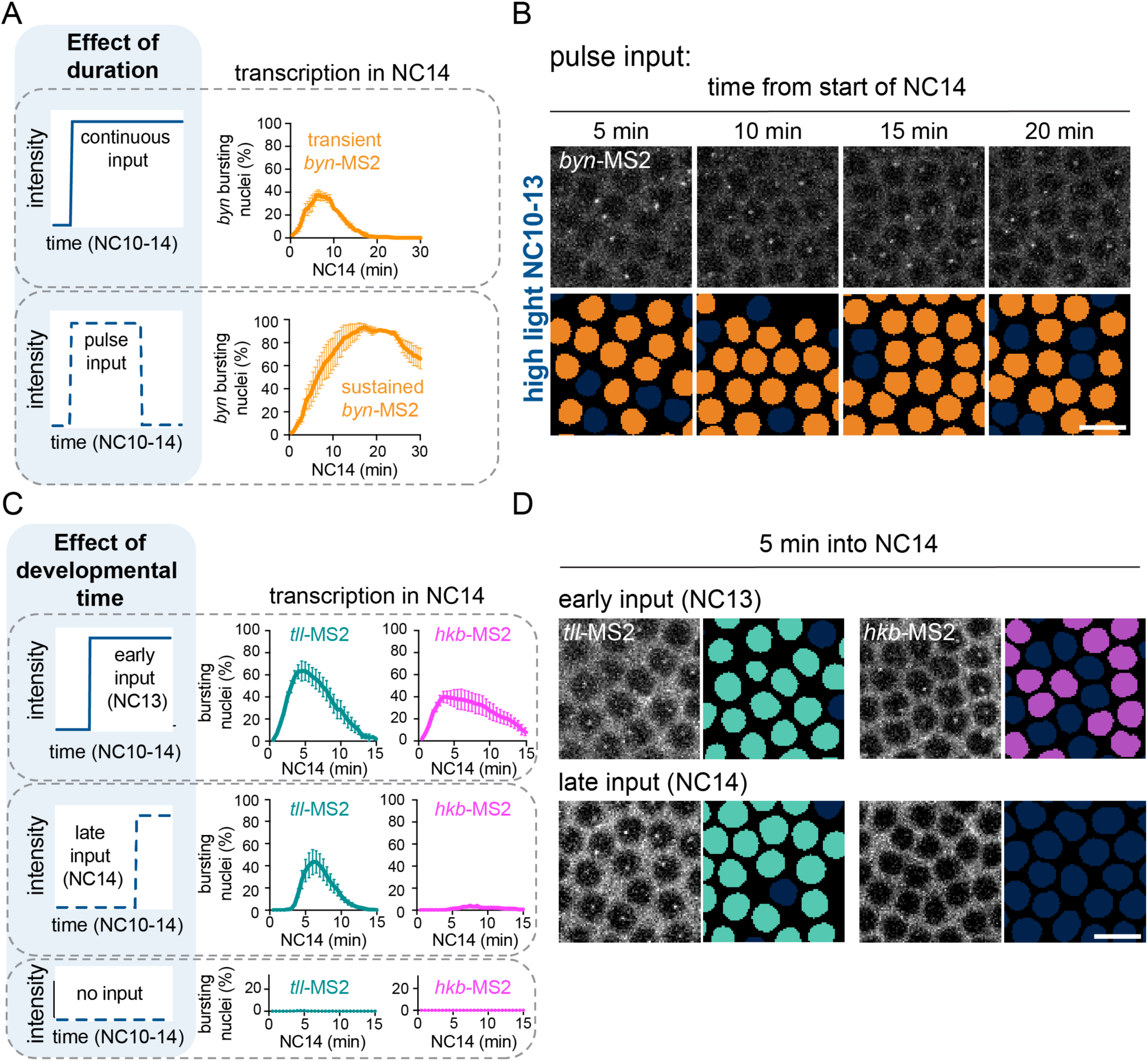
Dynamic manipulations confirm the conceptual model of the *byn* circuit. **(A)** Left: Schematic showing two inputs with different durations: (top left) a long, continuous duration from NC10-14 and (bottom left) a shorter, pulsed duration from NC10-13. Right: Graph showing proportion of nuclei in a *byn* bursting state during NC14 under these conditions. The top right graph is taken from the same dataset as Figure 1F. Mean ± SEM, n = 3-4 embryos. **(B)** *byn*-MS2 bursting during NC14 upon a pulse of high light stimulation that ends in NC13. Orange nuclear masks show nuclei with a burst in corresponding frames. Scale bar is 10 µm. **(C)** Left: Schematics show high inputs varied in their start time (NC13, NC14, or no input). Right: Graphs show the proportion of nuclei in a *tll* (teal) or *hkb* (magenta) bursting state during NC14 under these conditions. Mean ± SEM, n = 3-5 embryos in each condition. **(D)** *tll-* and *hkb*-MS2 bursting at 5 min into NC14 after the light begins at the start of NC13 (top) or NC14 (bottom). Nuclear masks (*tll* in teal, *hkb* in magenta) show nuclei with a burst in corresponding frames. Scale bar is 10 µm.

One key feature of our conceptual model is a time delay before *hkb* expression that depends on ERK stimulus intensity. However, an alternative model could hold that the timing of *hkb* transcription is primarily controlled by developmentally-regulated changes in its sensitivity to ERK. For example, *hkb* transcription may exhibit different sensitivities at different nuclear cycles, switching from being resistant to low-dose ERK stimulation during NC11 to being more sensitive during NC13-14. We reasoned that optogenetics could also be used to distinguish these models. By applying the same light input in different nuclear cycles and measuring the timing of *tll* and *hkb* expression, it is possible to separate a stimulus-response delay from a developmental timer.

We illuminated OptoSOS/MCP embryos using the high light condition at two different time points – an “early” stimulus beginning in NC13 or a “late” stimulus beginning at the start of NC14 (**Figure 3C**, left). We then assessed transcription of both *tll* and *hkb* using endogenous MS2 reporters. For *tll*, we observed a robust, immediate pulse of transcription in NC14, regardless of whether the light began in NC13 or 14 (**Figure 3C-D**, left, **Movie S4**). Even light stimulation in early NC14 produced an extremely rapid *tll* response, with MS2 foci detectable within just 4 minutes of illumination (**Figure 3C**-center panel) in agreement with previous findings (Keenan et al. 2020). Taken together with our observation of transcription in NC11 for stimuli delivered in NC10 (**Figure 2**), these data suggest that *tll* remains highly sensitive to ERK activity from NC10 to early NC14. For *hkb*, we observed that bright illumination in NC13 was sufficient to drive a pulse of transcription in NC14, but not when the illumination began in NC14 (**Figure 3C-D**, right, **Movie S4**). In sum, *tll* is always induced immediately following ERK activation, whereas *hkb* exhibits delayed transcription, regardless of the developmental time at which the stimulus is applied. Taken together with our observation of a dose-dependent delay in *hkb* expression, these results support a model where the dynamics of the *tll/hkb/byn* system are intrinsic to this incoherent feedforward circuit, and not externally imposed by developmental time. More broadly, the experiments described above demonstrate the power of optogenetic manipulations to interrogate candidate models of developmental signal interpretation.

### Non-local responses to localized Ras activation regulate *brachyenteron* spatial patterning

We have thus far investigated the temporal dynamics of *byn* expression, but prior work also suggests that its spatial pattern may be under complex regulation. In mutant embryos lacking the endogenous terminal pattern, an all-or-none OptoSOS input at the poles is sufficient for normal embryogenesis even though it eliminates both quantitative differences in Ras activity and spatial differences between *tll* and *hkb* expression (Johnson et al. 2020). OptoSOS embryos were also viable when all-or-none light inputs were applied on top of the endogenous gradient (Johnson et al. 2017). These results pose a conundrum because uniform high ERK activity and a complete overlap between *tll* and *hkb* expression would be naively predicted to eliminate a stripe of sustained *byn* expression. Having shown that *byn* dynamics might also be regulated by the temporal patterns of *tll* and *hkb* transcription, we next performed a closer examination of the circuit’s dynamics near the spatial boundary of an all-or-none OptoSOS stimulus (**Figure 4A**).

**Figure 4:**
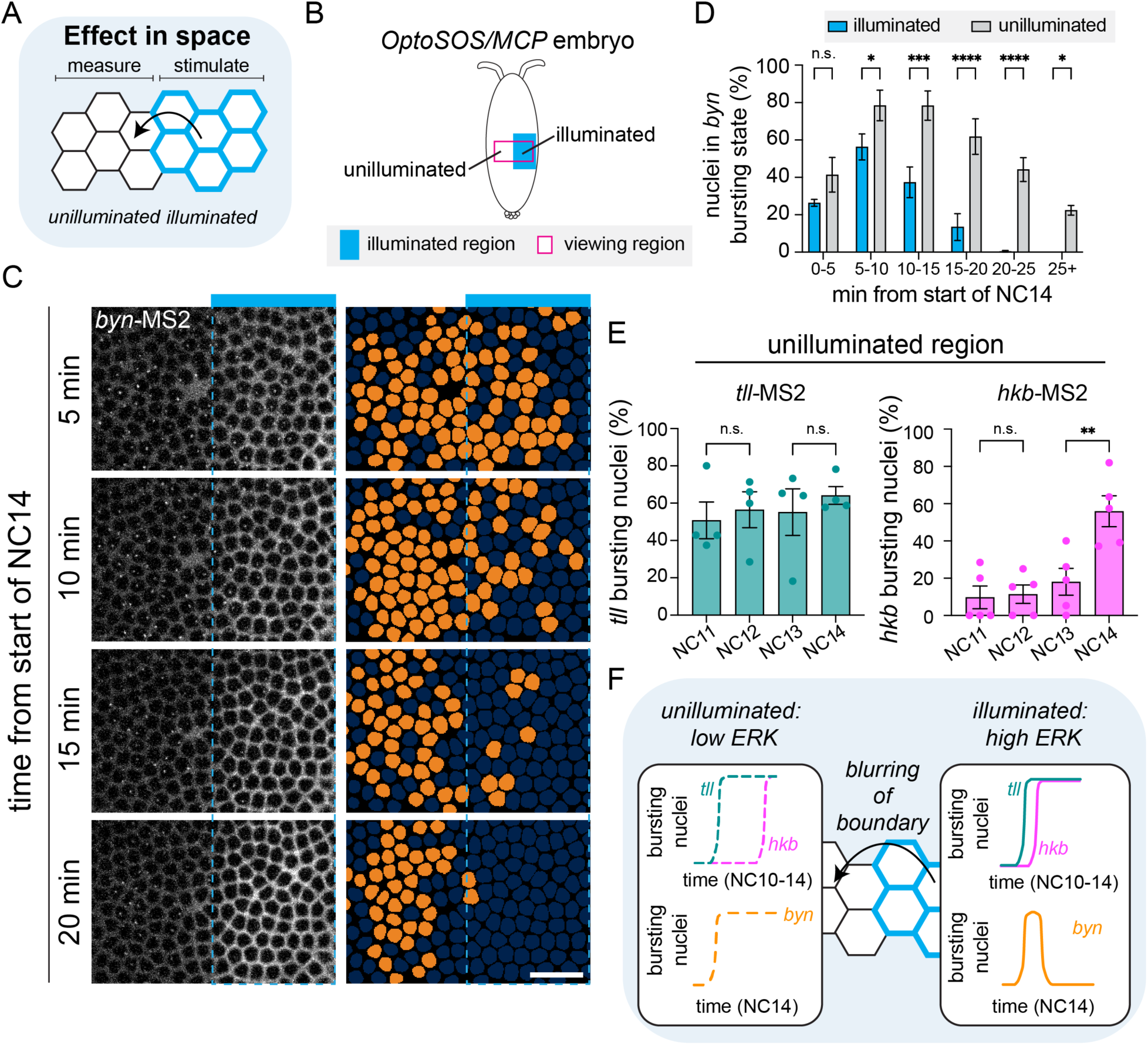
An all-or-none OptoSOS input produces a stripe of *byn* expression. **(A)** Schematic showing the potential for non-autonomous effects in space. The illuminated region is marked by blue membrane, but there may be effects in the unilluminated region. **(B)** Schematic showing how the ventral viewing window (magenta) extends into the unilluminated region at the same anterior posterior position. **(C)** *byn*-MS2 bursts in NC14 upon high illumination. The blue bar shows the illuminated portion of the viewing window, and the dashed blue line shows the illumination boundary. Orange nuclear masks show nuclei with a burst in corresponding frames. Scale bar is 10 µm. **(D)** Proportion of nuclei in the *byn* bursting state through NC14. Significantly more nuclei remain in the bursting state in the unilluminated region in later NC14. Mean ± SEM, n = 4 embryos in each condition. **(E)** Proportion of nuclei in the bursting state in each nuclear cycle for *tll* and *hkb* in the unilluminated region. Each point represents one embryo. Mean ± SEM, n = 4 embryos for each condition. Significance for all graphs derived from ANOVA with Sidak’s post hoc test. **(F)** Schematic for how temporal dynamics and blurring together contribute to formation of a *byn* stripe by an all-or-none input. In the illuminated region, ERK is high, *hkb* expression is only slightly delayed and *byn* is transient. In the unilluminated region, there is a lower level of ERK activity, the delay in *hkb* expression is large, and *byn* is sustained.

To quantify *tll, hkb,* and *byn* expression near the illumination boundary, we simply extended our analysis from prior experiments into the unilluminated half of each embryo (**Figure 4B**, **Figure S4A-B**). Our viewing window comprised both illuminated and unilluminated regions at the same anterior-posterior position and was symmetric across the ventral axis, thus enabling comparisons between these regions without introducing anterior-posterior or dorsoventral bias. Remarkably, when we considered the illuminated and unilluminated regions together under high light, we now observed the dynamic formation of a *byn* stripe in response to the all-or-none input (**Figure 4C**, **Movie S5**). Initially, *byn*-MS2 foci were detected throughout both the illuminated and unilluminated region, up to 50 μm away from the illumination boundary. After about 15 minutes, *byn* foci were lost in the illuminated region but persisted in the unilluminated region (**Figure 4C-D**). This non-local effect only occurs under high light; under low light there is no *byn* expression in the unilluminated region (**Figure S4C-E**). In sum, a bright all-or-none light pattern produced a *byn* stripe in the nearby unilluminated region with similar spatial dynamics to the wild-type pattern but rotated 90 degrees to match the illumination boundary.

This observation of sustained *byn* expression in the unilluminated region suggested that this region may also exhibit rapid *tll* expression but delayed or absent *hkb* expression. To test this prediction, we analyzed the unilluminated regions of *tll-*MS2 and *hkb-*MS2 embryos subjected to the high light condition. This analysis revealed that *tll* was transcribed beginning in NC11, whereas *hkb* transcription was again delayed until NC14 (**Figure 4E**). These results suggest that for *tll, hkb,* and *byn*, the unilluminated region of embryos subjected to a high light stimulus is functionally equivalent to the illuminated region under low light conditions (**Figure 4F**).

How does a uniform optogenetic input to the Ras/ERK pathway act at a distance to induce *tll*, *hkb*, and *byn* transcription ~50 μm outside of the illuminated region? We envisioned two possibilities. First, light scattering or imprecision of the optogenetic input might lead to blurring of the all-or-none pattern to accidentally form a gradient. Second, active cellular components might diffuse within the syncytial embryo so that an initially sharp stimulus pattern would produce a graded response. We took two strategies to discriminate between these possibilities. First, we quantified the spatial responses at multiple nodes including our light input, OptoSOS membrane recruitment, an ERK activity biosensor (Moreno et al. 2019), and Tll/Hkb protein distributions (**Figure S5A-D**). Overall, we found that our light stimulus and membrane OptoSOS recruitment formed relatively precise boundaries, whereas ERK activity and Tll/Hkb expression were graded and extended ~50 μm away from the illumination boundary. Second, we tested for intracellular component mixing using a photoconvertible tdEOS-tubulin construct (Lu et al. 2013) whose green-to-red photoconversion we could control using the same optical path. Our illumination setup was able to produce an initially-sharp pattern of photoconverted tdEOS that subsequently underwent substantial blurring over a time period of 10 min; no blurring occurred when the same embryos were photoconverted after cellularization (**Figure S5E-F**). Taken together, these data suggest that an all-or-none Ras stimulus is converted into a graded transcriptional response at least in part through the diffusion or transport of activated intracellular components.

### All-or-none Ras activation at the posterior pole rescues *brachyenteron* stripe formation

As a final test of our model, we asked if the same uniform optoSOS input could rescue patterning of the *byn* stripe at the endogenous position in an embryo lacking the endogenous terminal pattern. Embryos mutant for *torso-like (tsl)*, an activator of the Trunk ligand, lack terminal ERK signaling and do not express *byn* (**Figure 5A**, left plot) (Savant-Bhonsale and Montell 1993). Although we recently reported that an all-or-none optogenetic stimulus at the poles was sufficient to rescue subsequent development (Johnson et al. 2020), we did not examine *byn* expression under these conditions.

**Figure 5:**
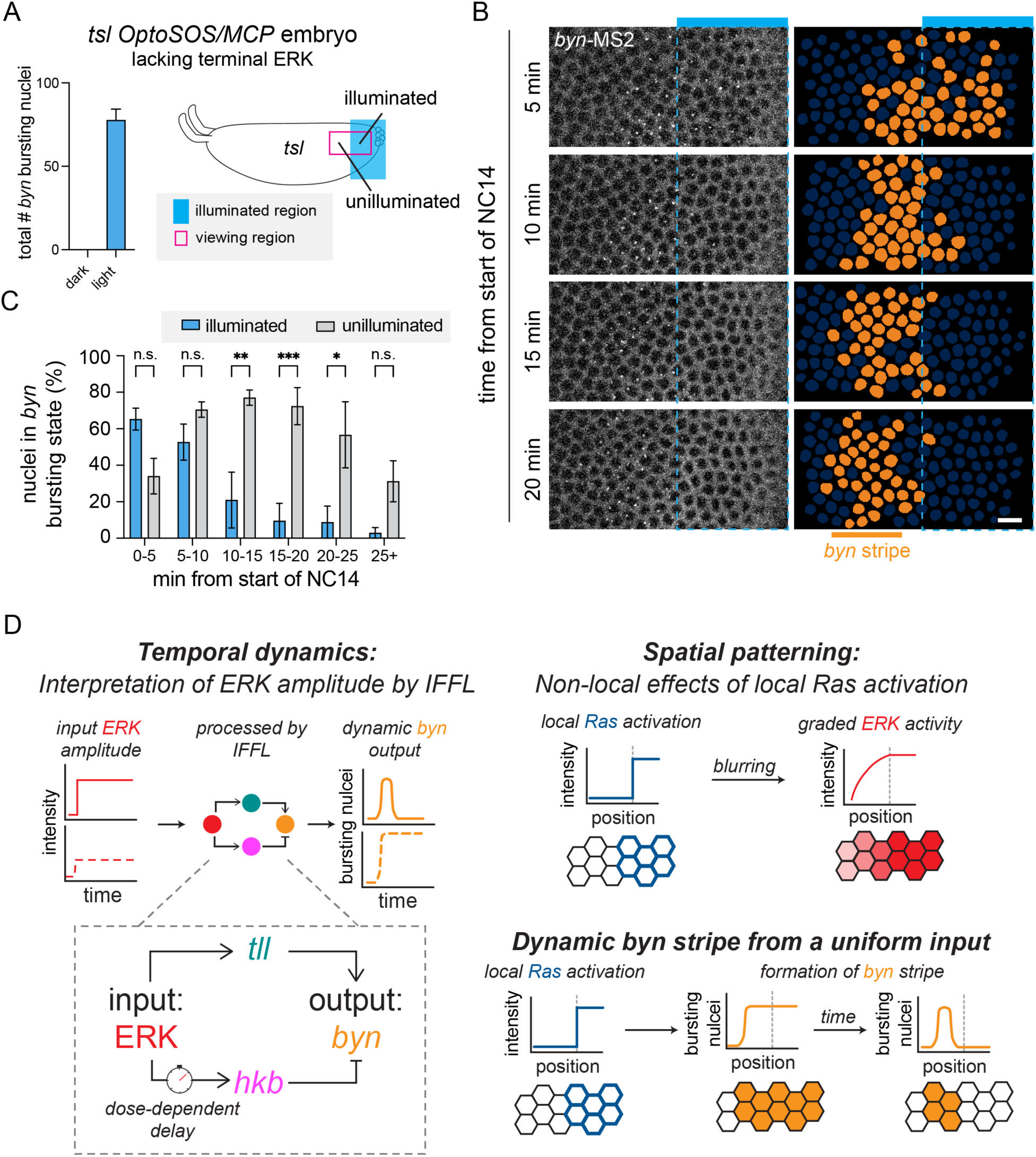
A uniform high input at the pole of a *tsl* mutant embryo produces a stripe of *byn*. **(A)** Left: Plot of total number of nuclei with a *byn* burst in the viewing region of a *tsl* OptSOS/MCP embryo under dark and light conditions. Mean ± SEM, n = 3-4. Right: Schematic showing the illumination region at the posterior pole of a *tsl* mutant OptoSOS/MCP embryo. The viewing window (magenta) extends into the unilluminated region. **(B)** *byn*-MS2 bursts in NC14 upon high illumination. The blue bar shows the illuminated portion of the viewing window, and the dashed blue line shows the illumination boundary. Orange nuclear masks show nuclei with a burst in corresponding frames. Scale bar is 10 µm. **(C)** Proportion of nuclei in the *byn* bursting state through NC14. Significantly more nuclei remain in the bursting state in the unilluminated region in later NC14. Mean ± SEM, n = 4 embryos in each condition. Significance from two-way ANOVA with Sidak’s post hoc test. **(D)** Conceptual model of *byn* stripe formation that incorporates temporal dynamics and spatial patterning. Left: ERK inputs are processed by *tll* and *hkb* in an incoherent feedforward loop architecture to produce *byn* outputs with different dynamics. Right: Sharp Ras inputs are blurred to produce graded ERK activity in space. Together, these processes explain how an all-or-none input produces a *byn* stripe.

We generated OptoSOS/MCP embryos from mothers that were also homozygous for the *tsl* loss-of-function mutation (**Figure 5A**, right schematic). Unlike in prior experiments, where we restricted illumination to the ventral surface of the embryo away from the poles, here we applied our light stimulus directly at the posterior pole beginning in NC10 and visualized *byn*-MS2 transcription in both the illuminated and neighboring unilluminated regions. At early time points in NC13 (**Figure 5B**, 5 min) we observed *byn* expression in both the illuminated and unilluminated regions. However, by 10 min into NC14, *byn* resolved into a stripe in the unilluminated region that persisted for at least 20 min (**Figure 5B-C**). These data demonstrate that *byn* stripe formation in response to a simple all-or-none stimulus is a robust property of its regulatory circuit that can occur both at endogenous and ectopic positions in the embryo.

## Discussion

Here we have dissected the regulation of the *byn* stripe by combining precise optogenetic inputs in space and time with live biosensors of target gene expression. Using ectopic activation of Ras on the ventral side of wild-type embryos, we defined high- and low-amplitude OptoSOS inputs that induce distinct *byn* transcriptional dynamics – a pulse of expression in early NC14 versus more sustained expression – that match its endogenous responses in the posterior terminus and stripe-forming region. We then used these conditions to characterize the *tll* and *hkb* inputs that explain these *byn* dynamics in space and time.

Our approach yielded novel insights about both the temporal and spatial interpretation of ERK inputs to pattern the *byn* stripe. First, differences in signal amplitude are interpreted through the timing of *tll* and *hkb* expression (**Figure 5D**, left). The onset of *tll* expression is always rapid, occurring in as few as 4 minutes after signaling onset, whereas there is a dose-dependent delay in the onset of *hkb* expression. This delay in *hkb* expression is a function of Ras/ERK input amplitude, not of developmental time. These data are consistent with previous observations in OptoSOS embryos that *hkb* RNA only accumulates to high levels in response to blue light inputs over 30 min (Johnson and Toettcher 2019). They also broaden our conception of the thresholds for *tll* and *hkb* expression*: tll* and *hkb* can be induced by inputs of the same amplitude, but *hkb* requires that the signal persist for a longer time. If the amplitude is low enough, the signal must persist longer than the developmental window allows, and *hkb* is never expressed. Thus, cumulative dose of ERK input (amplitude integrated over time) appears to be the relevant feature sensed by the circuit, as has been proposed for the terminal pattern as well as other systems (Johnson and Toettcher 2019; Gillies et al. 2017). The circuit then processes this input through the relative timing of *tll* and *hkb*, rather than simply their presence, to determine local *byn* expression.

This more nuanced understanding of *byn* regulation resolves a conundrum of the endogenous pattern: how can *byn*’s transient pulse of expression in the high-ERK, Hkb-positive domain be reconciled with the presence of its inhibitor? Here we show that at the high light levels which produce a comparable pulse of *byn* transcription, *hkb* transcription is delayed relative to *tll* and this delay is also evident in the accumulation of their protein products. Thus, there is a temporal window where only the positive regulator is present, allowing for a pulse of *byn* expression, prior to accumulation of the repressor. This sequential appearance of Tll and Hkb was hypothesized during the initial characterizations of posterior patterning (Casanova 1990) but has only now been directly shown. It is interesting to note that Tll has been characterized as a transcriptional repressor, implying that there is an intermediate node between *tll* and *byn* (Morán and Jiménez 2006). However, the identity of this node and how it affects the timing of *byn* activation and repression remain unknown.

Our improved understanding of *byn* regulation also explains how a *byn* stripe can form in conditions where *tll* and *hkb* have the same spatial domain, as in the rescue experiment of Johnson et al (2020). Our current study revisits these results with improved tools, in particular endogenously-tagged transcriptional reporters of *tll* and *hkb* that are able to clearly resolve differences in transcriptional dynamics that were obscured by enhancer-based reporter constructs. We find that stimulus conditions that support sustained *byn* can also support *hkb* expression in NC14, but under these conditions *hkb* expression is largely absent from earlier nuclear cycles. This observation differs from the wild-type pattern, where the *byn* stripe forms in a Tll-only region that is presumably formed from an ERK gradient that attains even lower activity levels than our optogenetic inputs. It is possible that the network dynamics we report here provides robustness to the *byn* circuit, allowing it to produce different outputs for even a narrow range of input strengths.

Our study reveals that the *tll*-*hkb*-*byn* circuit can be classified as an incoherent feedforward loop with rapid activation and delayed repression, a circuit with well-characterized pulse-generation and stripe-forming properties (**Figure 5D**, left) (Alon 2007; Schaerli et al. 2014). A unique feature of this circuit however is that the delay in *hkb* expression is dose-dependent, meaning that differences in signal amplitude are converted to differences in *hkb* dynamics and thus different *byn* responses (i.e., transient if *hkb* onset is fast, sustained if *hkb* onset is slow). Interestingly, a similar dose-dependent delay in transcriptional onset was recently shown for *T48* downstream of the Dorsal gradient (Carmon et al. 2021; Keller et al. 2020). What is the mechanism underlying this delay in *hkb* onset? The dose-dependence of *tll* and *hkb* has been a longstanding open question even without the complexity of temporal dynamics (Furriols and Casanova 2003; Smits and Shvartsman 2020). ERK signaling activates transcription of both *tll* and *hkb* through relief of the same repressor, Capicua (Cic), and it is unclear why these genes would require different doses of ERK signaling. Our experiments rule out a few possible explanations. Developmental time does not appear to be critical, given that the delay in *hkb* transcription is observed regardless of when light is applied and both the *tll* and *hkb* loci are known to be accessible early (Blythe and Wieschaus 2016). We can also rule out interactions with other components of the anterior-posterior patterning machinery given that we are able to produce an ectopic *byn* stripe rotated 90 degrees from its endogenous counterpart. One intriguing possibility, supported by previous ChIP-Seq results, is that Cic leaves *hkb’s* enhancers more slowly than those of *tll* (**Figure S6**) (Keenan et al. 2020). It is also possible that signaling dependent chromatin changes are involved (Semprich et al. 2022; Hannon et al. 2017). These models will be tested in future studies.

The second major finding is that the boundary of a uniform OptoSOS input is blurred in space downstream of Ras to produce two domains from a single input – a transient *byn* domain within the high-ERK illuminated region and a sustained domain in the low-ERK unilluminated region (**Figure 5D**, right). These non-local effects of a local Ras input are most likely mediated by diffusion of active intracellular components, a well-established contributor to developmental patterning in the syncytial *Drosophila* embryo (Driever and Nüsslein-Volhard 1988; Gregor et al. 2005). It remains unknown whether the endogenous terminal dpERK gradient is produced from a similar gradient of active Torso receptors, or is due to the combination of a discrete domain of Torso activity at the poles and cytoplasmic diffusion of downstream components (Casanova and Struhl 1989; Coppey et al. 2008). If the latter model is correct, the developmental rescue by an all-or-none OptoSOS input may not be an example of a simple input replacing the function of a complex one, but rather a good approximation of endogenous activation in the terminal system. A number of systems once thought to depend strictly on input concentration have similarly been shown to depend on an unexpectedly simple form of the input (Zhu et al. 2022; Economou et al. 2022).

Altogether, this work provides a blueprint for dissecting a developmental circuit with optogenetic tools to reveal new insights about network architecture. We have manipulated amplitude, duration, timing, and spatial pattern of the signal to understand the contributions of each factor to signal interpretation. This framework will be an effective strategy for dissecting other developmental circuits in the future.

## Material and Methods

### Drosophila melanogaster stocks

Fly stocks used in this paper include the maternal tubulin Gal4 driver *67; 15* (*P{matα-GAL-VP16}67; P{matα-GAL-VP16}15,* BDSC #80361), *UAS-OptoSOS* (Johnson et al. 2017), *UAS-OptoSOS; MCP-mCherry* (McFann et al. 2021), endogenous MS2-tagged *tll-24xMS2*, *hkb-24x MS2,* and *byn-24xMS2* (Keenan et al. 2022), *67; 15 tsl^4^*/*TM3 Sb* and *MCP-mCherry; UAS-OptoSOS tsl^4^/TM3 Sb* (Johnson et al. 2020), *tub-miniCic-NeonGreen* (a gift from Romain Levayer), *vasa-mCherry* (a gift of Hernan Garcia), *tll*-*mCherryLlamaTag* and *hkb-mCherryLlamaTag* (this paper), and *UAS-alphaTub84B.tdEOS* (BDSC #51313). Flies were cultured using standard methods at 25 °C.

### Embryo preparation

For all imaging experiments, a cage of parental flies was placed in the dark at room temperature for at least 2 days prior to collection with standard apple juice plates and yeast paste. For wild-type experiments, female *67/OptoSOS; 15/MCP-mCherry* virgins were mated with homozygous MS2 males. For *tsl* mutant experiments, female *67/MCP-mCherry; 15 tsl^4^/OptoSOS tsl^4^* virgins were mated with homozygous *byn*-MS2 males. Embryos were mounted between a semi-permeable membrane and a coverslip in a 3:1 mix of halocarbon 700/27 oil. Wild-type embryos were mounted with their ventral side facing the coverslip and *tsl* embryos were mounted with their lateral side facing the coverslip.

### Live microscopy and patterned optogenetic stimulation

All imaging was done at room temperature at 40x on a Nikon Eclipse Ti microscope with a Yokogawa CSU-X1 spinning disk and an iXon DU897 EMCCD camera. Optogenetic stimulation was performed with a Mightex Polygon digital micromirror device using an X-Cite LED 447-nm blue LED. Embryos with similar red fluorescence levels were selected for imaging with the goal of choosing embryos expressing similar levels of OptoSOS. For wild-type embryos, a stimulation region was drawn at a position slightly posterior to the center of the embryo encompassing only the right half of the embryo, with the ventral midline as the illumination boundary. For *tsl* embryos, a stimulation region was drawn encompassing roughly the most posterior 60 μm of the embryo. Unless otherwise noted, blue light stimulation began in NC10, when nuclei could first be observed at the surface. A 0.6 sec pulse of blue light was applied to the embryo every 30 sec. Light amplitude was varied by changing the LED power over an 8-fold range. Images were acquired every 30 sec with 1 μm z slices over a range of 22-30 μm. At high light intensities, SSPB-tagRFP-SOS^cat^ rapidly relocalizes to the membrane. Any embryos without visible SOS^cat^ localization were excluded. At low light intensities, this membrane enrichment is only barely detectable.

### Quantification of MS2 bursts

Images were processed using ImageJ. All images containing MS2 were background subtracted using rolling ball subtraction with radius 50 pixels. A maximum projection was then made with z slices containing the nuclei. Images were analyzed manually, and each nuclear cycle was analyzed independently. For each nucleus, the position at the start of each nuclear cycle was identified. To account for any variability in mitotic wave speed, the start of a nuclear cycle was defined as the time when the reformed nuclei were first visible within the quantified region.

The nucleus was then annotated with the time of the first and last visible MS2 focus, defining the “bursting duration”. The nucleus was defined to be in a “bursting state” at any timepoint within this bursting duration, even if an MS2 focus is not detected. The proportion of nuclei in a bursting state during each nuclear cycle or at a given timepoint within a nuclear cycle was calculated out of the total number of nuclei. To determine the “time to 50% nuclei in bursting state” for each embryo, we identified the first nuclear cycle in which at least 50% of nuclei were in a bursting state and at least 50% of nuclei remained in a bursting state for all subsequent nuclear cycles. We then identified the timepoint (measured in minutes since illumination began) at which 50% of nuclei had entered the bursting state within that nuclear cycle.

We observed some heterogeneity within the viewing region. At low light intensities, *tll* and *byn* varied in how far in the anterior direction their transcription extended in the illuminated region. To account for this variation, we quantified a box of defined size (50 x 45 μm) positioned at the most anterior extent of transcription in NC14 when comparing high and low light (See **Figure S2A-B**). Similarly, when comparing transcription under high light in the illuminated and unilluminated region, we found that *tll*, *hkb*, and *byn* all varied in how far from the illumination boundary their transcription extended. Therefore, we similarly defined a 30 x 75 μm box that shifted in the unilluminated region to a position at the farthest extent of transcription away from the illumination boundary in NC14 (See **Figure S4A-B**). Any nuclei within this quantification box were included in quantifications.

### Quantification of OptoSOS membrane enrichment

To measure membrane enrichment of OptoSOS, we segmented out the nuclei from the background subtracted and maximum projected images, leaving only the cytoplasm/membrane portions. In NC14, when this analysis was done, there is very little visible cytoplasm in these planes so we included any non-nuclear region in the membrane analysis. To measure OptoSOS recruitment as a proxy for ERK signaling (**Figure 2D**), we measured the difference in mean membrane intensity in the illuminated region and an unilluminated region at the same anterior-posterior position. To measure the profile of OptoSOS recruitment relative to the illumination boundary (**Figure S6B**), we measured the median intensity at each pixel position at increasing distances from the boundary and normalized it to the highest intensity measurement in each embryo.

### Burst intensity visualizations

Nuclei were segmented using a model fine-tuned from Cellpose’s “cyto2” model, trained using default hyperparameters on manually segmented frames. MS2 bursts were identified through manual segmentation. To produce burst intensity visualizations, the nucleus mask nearest to each identified burst was colored to indicate detection.

### Statistical analysis

All statistical analysis was performed in Prism 9 (Graphpad). Details for statistical tests used can be found in the figure legends. Figure legends also indicate the number of embryos or nuclei analyzed for each condition (n). All graphs show the mean and SEM. In all cases, significance was defined as *p-value < 0.05, **p-value < 0.01, ***p-value < 0.001, and ****p-value < 0.0001. n.s. indicates no significance.

## Acknowledgements

We thank Romain Levayer for the kind gift of *miniCic-NeonGreen* flies and Hernan Garcia for the kind gift of *vasa-mcherry* flies. We thank Eric Wieschaus and all members of the Toettcher and Shvartsman labs for helpful discussions. This work was supported by the Hertz Fellowship and the NSF Graduate Research Fellowship Program (S.E.M), NIH grant F32GM119297 (H.E.J), NIH grants R01HD085870 and R01GM134204 (S.Y.S.), and NSF CAREER Award 1750663 and NIH Grant U01DK127429 (J.E.T.). Stocks obtained from the Bloomington Drosophila Stock Center (NIH P40OD018537) were used in this study.

## Author contributions

E.K.H., S.E.M., S.Y.S., and J.E.T. designed the study. E.K.H. performed experiments. S.E.M. and H.E.J. performed preliminary experiments. E.K.H. and H.R.O. analyzed the data and generated the figures. L.Y. generated the LlamaTag flies. E.K.H. and J.E.T. wrote the manuscript with feedback from all authors.

