## Supplemental Materials for "Dynamics of an incoherent feedforward loop drive ERK-dependent pattern formation in the early *Drosophila* embryo"

Supplemental Figures S1-S6

Supplemental Figure S1 (related to Figure 1)
Supplemental Figure S2 (related to Figure 1 and 2)
Supplemental Figure S3 (related to Figure 2)
Supplemental Figure S4 (related to Figure 4)
Supplemental Figure S5 (related to Figure 4)
Supplemental Figure S6 (related to discussion)

Supplemental Materials and Methods

Supplemental Table S1 (related to Supplemental Materials and Methods)

Legends for Supplemental Movies S1-S5

Supplemental Movie S1 (related to Figure 1)
Supplemental Movie S2 (related to Figure 2)
Supplemental Movie S3 (related to Figure 3)
Supplemental Movie S4 (related to Figure 3)
Supplemental Movie S5 (related to Figure 4)

Supplemental References

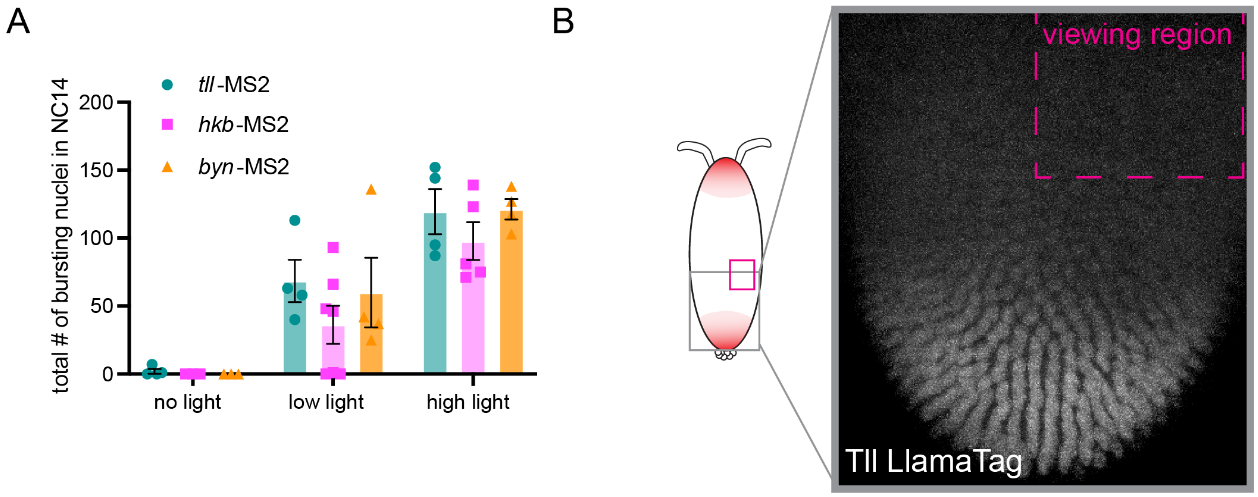


**Figure S1: The ventral region used in this study does not express endogenous *tll*, *hkb*, or *byn*** **(A)** Plot of total number of nuclei that burst in response to high, low, and no light for *tll*, *hkb*, and *byn*. There is no transcription of any of the targets under no light in this ventral region, indicating that all bursts we observe are induced by OptoSOS. Mean ± SEM, n = 3-7 embryos. **(B)** Image of the posterior pole of an embryo in the dark expressing Tll LlamaTag (see **Figure S3**). The viewing region used for all experiments is marked in magenta and is clearly distant from the region of endogenous Tll.

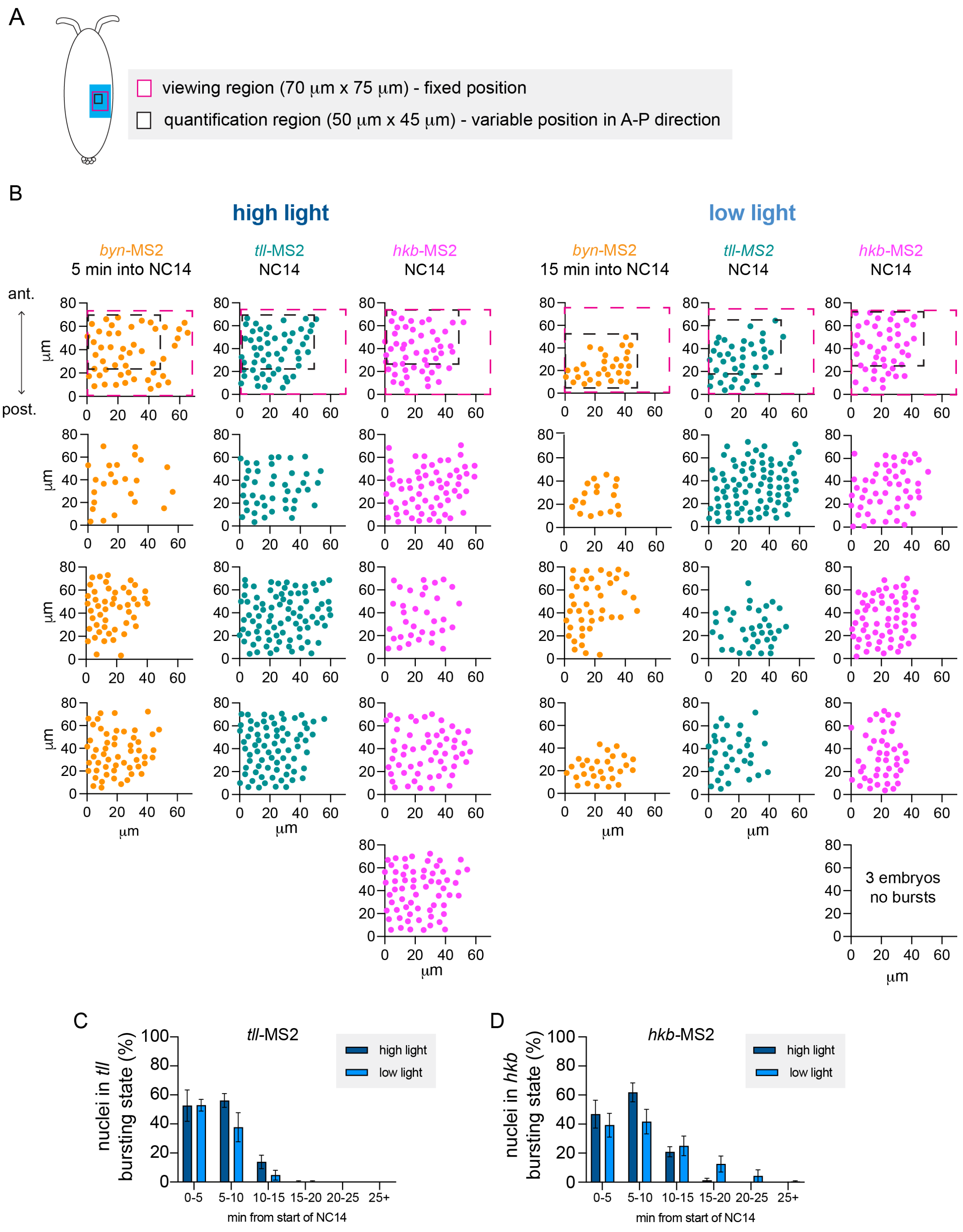
A
viewing region (70 μm x 75 μm) - fixed position
quantification region (50 μm x 45 μm) - variable position in A-P direction
B
high light
low light
byn-MS2
5 min into NC14
tll-MS2
NC14
hkb-MS2
NC14
byn-MS2
15 min into NC14
tll-MS2
NC14
hkb-MS2
NC14
ant.
post.
μm
3 embryos
no bursts
C
tll-MS2
nuclei in tll bursting state (%)
high light
low light
min from start of NC14
0-5
5-10
10-15
15-20
20-25
25+
D
hkb-MS2
nuclei in hkb bursting state (%)
high light
low light
min from start of NC14

**Figure S2: Characterizing *byn*, *tll* and *hkb* transcriptional response to high and low light**
We noticed that there was some variability in the expression domains of *byn, tll*, and *hkb* between embryos, with the most notable variation seen for the anterior extent of *byn* and *tll* expression under low light. **(A)** To control for this heterogeneity, we quantified only a 50 x 45 μm box at the most anterior portion of expression. *byn* dynamics in this region were similar between replicates. **(B)** Plots show variability in *byn* (orange)*, tll* (teal), and *hkb* (magenta) expression between embryos. Each plot represents the viewing region (magenta box) of one embryo and each point represents a nucleus that is in the bursting state at the specified timepoint. The chosen quantification region is shown for a few representative embryos (black box). **(C-D)** Proportion of nuclei in the *tll* (C) and *hkb* (D) bursting state through NC14. For the *hkb* low light condition, only embryos with bursts present were included. Mean ± SEM, n = 4-5 embryos in each condition.

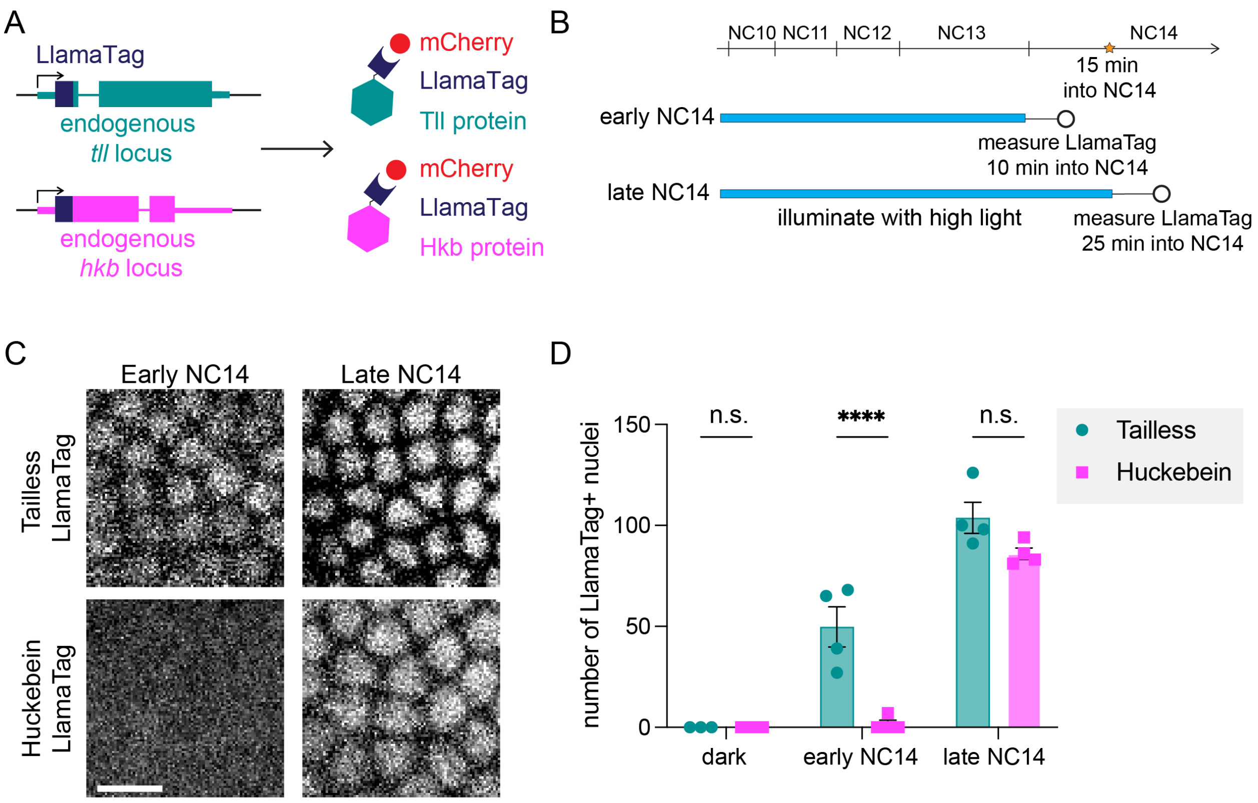


**Figure S3: Accumulation of Hkb protein is delayed relative to Tll protein**

**(A)** CRISPR/Cas9-tagged *tll* and *hkb* alleles with an anti-mCherry nanobody (LlamaTag) at the N-terminus. Upon translation of Tll or Hkb protein, the nanobody (dark blue) binds pre-folded mCherry protein (red) in the embryo, leading to accumulation of mCherry-bound Tll/Hkb protein in the nucleus. We generated embryos that expressed the OptoSOS system and mCherry (OptoSOS/mCherry embryos) and which also contained either the Tll-LlamaTag or Hkb-LlamaTag. **(B)** Schematic showing that illumination with high light begins in NC10 and continues until either 10 min into NC14 ("early") or 25 min into NC14 ("late"). The star shows the point 15 min into NC14 where transient *byn* transcription has ended. **(C)** Images show Tll or Hkb LlamaTag at early and late NC14. Nuclear localization indicates the protein is present. Scale bar is 10 µm. **(D)** Number of LlamaTag+ nuclei in each condition. Each dot represents one embryo. There are significantly more Tll+ nuclei in early NC14 but not in late NC14 by two-way ANOVA and Sidak's post hoc test. Mean ± SEM, n = 4 embryos.

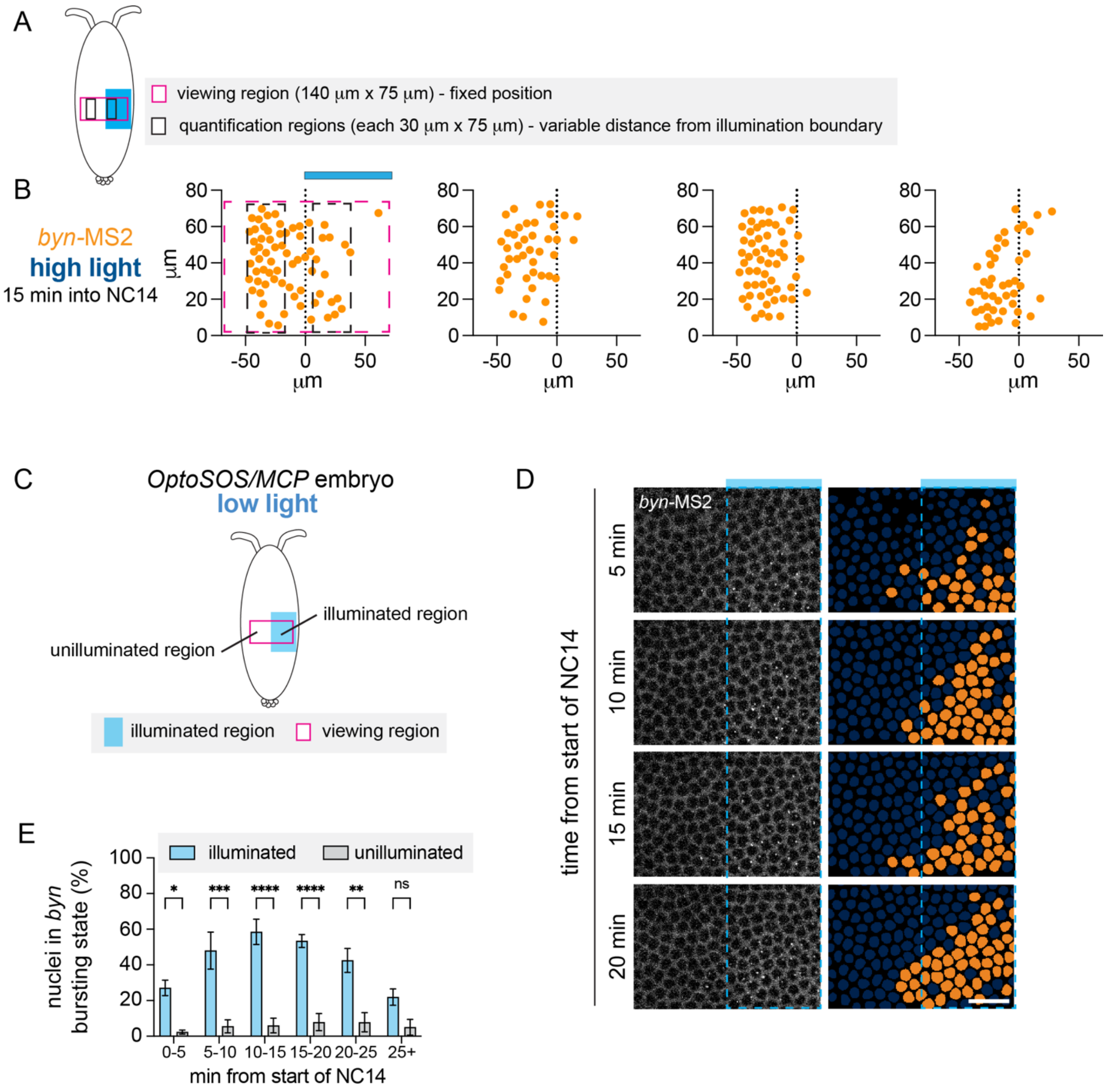


**Figure S4: Characterizing expression in the unilluminated region**

When comparing the illuminated and unilluminated regions, we noticed that there was variability in the position where the *byn* stripe forms relative to the illumination boundary. **(A)** To control for this heterogeneity, we quantified only a 30 x 70 µm box positioned at the furthest extent of *byn* expression. In the illumination region, the quantification region was fixed 5 µm away from the illumination boundary. **(B)** Plots show the variability in *byn* (orange) expression between embryos 15 minutes into NC14. At this timepoint, only nuclei expressing sustained *byn* are bursting. Each plot represents the viewing region (magenta box) of one embryo and each point represents a nucleus that is in the bursting state. The quantification region is shown for one representative embryo (black box). **(C)** We also measured expression of *byn* in the unilluminated region under low light. There was very little expression in the unilluminated region, consistent with ERK levels too low to induce expression. **(D)** Upon low light illumination, *byn*-MS2 bursts in NC14 were sustained in the illuminated region and there were very few bursts in the unilluminated region. The blue bar shows the illuminated portion of the viewing window, and the dashed blue line shows the illumination boundary. Orange nuclear masks show nuclei with a burst in corresponding frames. Scale bar is 10 µm. **(E)** Quantification shows the proportion of nuclei in the *byn* bursting state through NC14 upon low light stimulation. Mean ± SEM, n = 4 embryos in each condition. Significance from two-way ANOVA with Sidak's post hoc test.

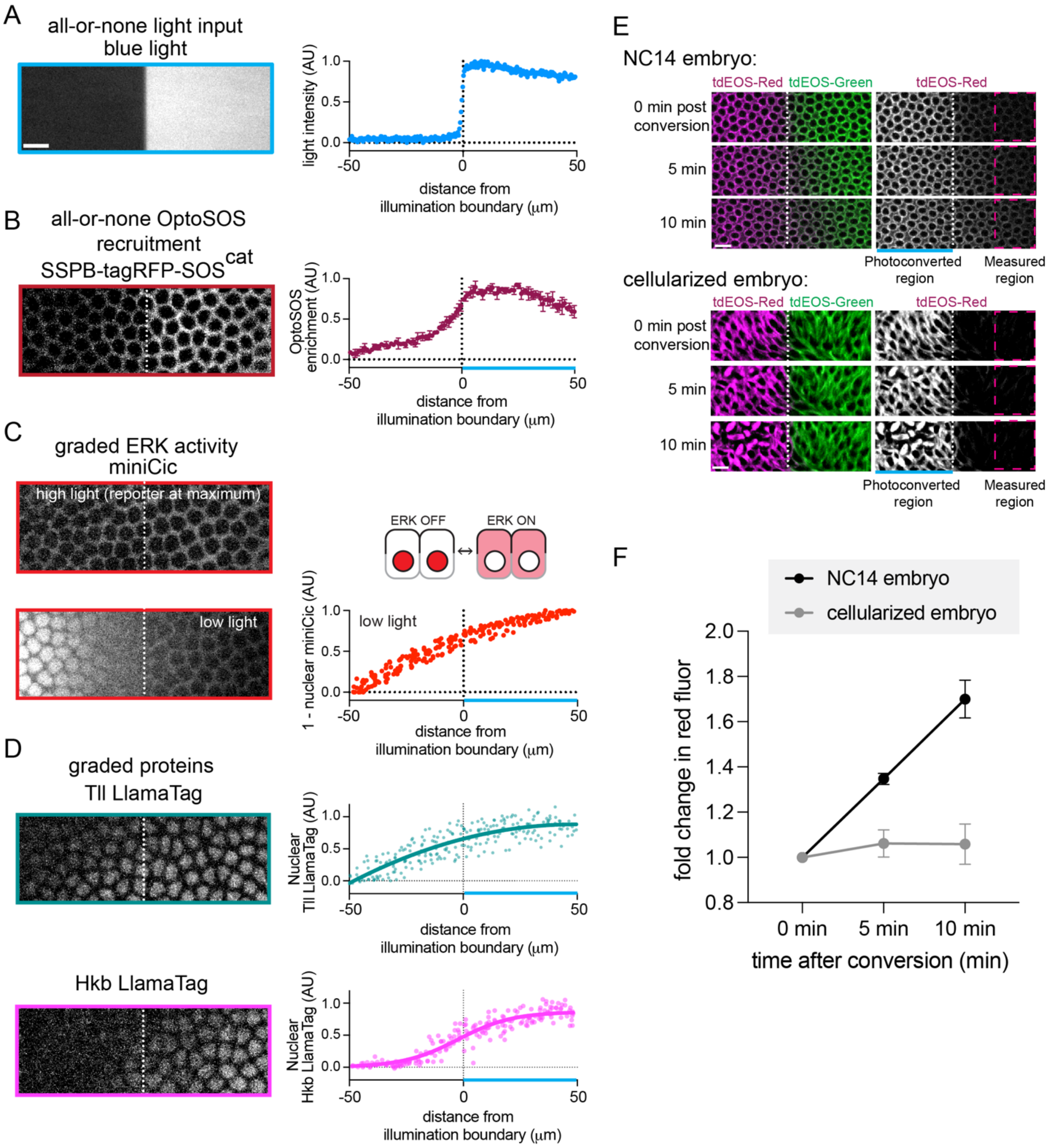


**Figure S5: Local Ras activation produces a diffusive gradient of ERK activation and gene expression**

We quantified the response of multiple effectors to our high light input on the ventral side. Light was applied from NC10 to NC14 and all measurements were taken in NC14. For each effector a representative image is shown on the left and quantification is shown on the right. The dotted vertical line represents the illumination boundary. Scale bars are 10 μm. **(A)** Blue light input forms a sharp boundary. Graph shows the intensity at each position from one representative image. **(B)**

Under high light, the SSPB-tagRFP-$SOS^{cat}$ component of OptoSOS is recruited to the membrane only in the illuminated region, with a sharp drop-off at the illumination boundary. Graph shows mean ± SEM intensity at each position from 3 embryos under high light. **(C)** We used miniCic, a reporter of ERK kinase activity, to assess ERK activity (Moreno et al. 2019). When ERK is off, miniCic is nuclear, and when ERK is on, miniCic relocalizes to the cytoplasm. Under high light, miniCic is cytoplasmic in both the illuminated and unilluminated regions, showing that ERK activity extends into the unilluminated region (upper image). The miniCic reporter saturates at relatively low levels of ERK, and so to understand the shape of the ERK gradient, we also applied low light. Under low light, there is a graded decrease in ERK activity at increasing distances from the illumination boundary (lower image, graph). Thus, a sharp boundary of OptoSOS activity is blurred into a gradient of ERK activity. Graph shows (1 - nuclear miniCic intensity) for n=3 embryos, each dot is one nucleus. **(D)** In late NC14, nuclear Tll LlamaTag (top) forms a shallow gradient, indicating that Tll protein is present up to 50 µm away from the illumination boundary. Nuclear Hkb LlamaTag (bottom) forms a steeper gradient. Graphs show nuclear LlamaTag intensity for n=3 embryos, each dot is one nucleus. The lines show the quadratic (Tll) and logistic (Hkb) function that best fits the data. **(E)** Images show tdEOS-tubulin at 0, 5, and 10 minutes post conversion from green to red in an early NC14 (top) and cellularized (bottom) embryo. The photoconverted region is to the left of the dotted white line and the dashed magenta box shows the quantified region 25 to 50 µm away from the illumination boundary. **(F)** Fold change in tdEOS-Red in the region 25 to 50 µm away from the illumination boundary in NC14 and cellularized embryos. n = 3 embryos for each condition. Mean ± SEM.

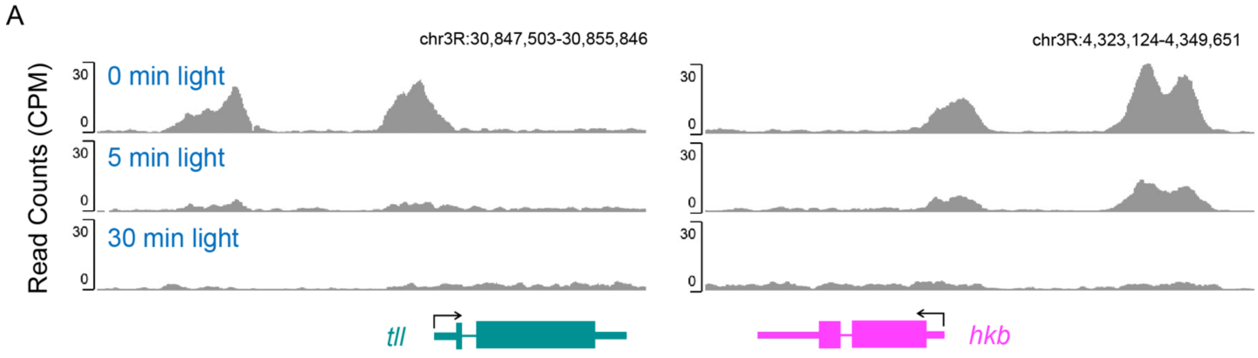


**Figure S6: Dynamics of Capicua loss after OptoSOS stimulation differ between *tll* and *hkb***
**(A)** ChIP-Seq profiles of Cic from Keenan et al (2020) were analyzed at the *tll* and *hkb* enhancers after OptoSOS stimulation for 0, 5, or 30 min. After 5 min, most Cic has been lost from the *tll* enhancer, but a peak remains at the *hkb* enhancer that is not lost until 30 min post stimulation.

### Supplemental Material and Methods

Generation of Tll and Hkb LlamaTag flies

Endogenous Tll-mCherryLlamaTag and Hkb-mCherryLlamaTag flies were generated using the scarless CRISPR/Cas9 editing method to insert the mCherryLlamaTag sequence into the N-terminus of Tll and Hkb (Bier et al. 2018; Bothma et al. 2018). In brief, homology arms were PCR amplified using TllN_left_Forward & TllN_left_Reverse, TllN_right_Forward & TllN_right_Reverse, HkbN_left_Forward & HkbN_left_Reverse, and HkbN_right_Forward & HkbN_right_Reverse primers from genomic DNA isolated from *nos-Cas9* flies (BDSC #78781). Then the homology arms were cloned into the pHD-mCherryLlamaTag-ScarlessDsRed vector using HiFi DNA Assembly (NEB #E5520) to generate pHD-Tll-mCherryLlamaTag and pHD-Hkb-mCherryLlamaTag. The pHD-mCherryLlamaTag-ScarlessDsRed vector was made by substituting the sfGFP sequence of pHD-sfGFP-ScarlessDsRed (Addgene #80811) with the mCherryLlamaTag sequence from Bothma et al. (2018). The CRISPR PAM site in the pHD-Hkb-mCherryLlamaTag plasmid were mutated by Q5 site-directed mutagenesis (NEB #E0554) using Hkb_Q5_Forward & Hkb_Q5_Reverse primers. The gRNA plasmids for Tll and Hkb were generated by Q5 site-directed mutagenesis PCR from pU6-BbsI-chiRNA vector (Addgene #45946) using TllN_gRNA_Forward & TllN_gRNA_Reverse and HkbN_gRNA_Forward & HkbN_gRNA_Reverse primers (Gratz et al. 2013). The homology arms and gRNA plasmids were co-injected into *nos-Cas9* flies by BestGene Inc. The dsRed cassette was subsequently removed by crossing to *nos-PBac* flies (Keenan et al. 2022). Primer sequences for cloning are listed in **Table S1**.

LlamaTag crosses and imaging

Female *67/vasa-mCherry; 15/ OptoSOS* virgins were mated with LlamaTag males. Embryos were stimulated with high light starting in NC10 on the right half of the ventral side. Under high light, bright membrane recruitment of SSPB-tagRFP-$SOS^{cat}$ obscured nuclear accumulation of the LlamaTags, and so we turned off the blue light for 10 minutes prior to imaging of LlamaTags. For the "early NC14" timepoint, the blue light was turned off when nuclei entered the $13^{th}$ mitosis and the LlamaTag was imaged 10 minutes later. For the "late NC14" timepoint, the blue light was turned off 15 minutes into NC14 and the LlamaTag was imaged 10 minutes later. For visualization of the gradient (**Figure S5D**), the late NC14 timepoint was used.

miniCic crosses and imaging

Embryos from *67/miniCIC-NeonGreen; 15/ OptoSOS* mothers were stimulated with high or low light starting in NC10 on the right half of the ventral side. Images were collected in early NC14.

Photoconversion crosses and imaging

Embryos from *67/UASp-alphaTub84B.tdEOS* ; *15/* + mothers were treated with bleach for 30 seconds to remove the eggshell, washed thoroughly with water, and mounted for imaging. Embryos at the start of NC14 or mid-gastrulation (roughly 4 hrs post egg lay) were selected. At one pole of the embryo, the entire field of view was stimulated for 1 minute with a 405 laser at 100% power. Immediately following the conversion, the stage was shifted to reveal the conversion boundary and roughly 5 x 1 µm z stacks were collected at 5-minute intervals.

Quantification of LlamaTags and miniCIC

All images containing LlamaTags were maximum projected with z slices containing the nuclei. Then, the background was subtracted using rolling ball subtraction with radius 50 pixels. For each embryo, the number of nuclei detectable by nuclear localization of the LlamaTag within the equal-sized viewing region was counted. To quantify the gradient of nuclear LlamaTag, a gaussian blur with radius 2 was applied to the background subtracted images. The intensity at roughly 100 points spanning the illuminated and unilluminated region was measured. These points were selected within nuclei if nuclei were detectable nearby. If there were no detectable nuclei, points were placed randomly. Then, all points within each embryo were normalized from 0 to 1. The gradient of nuclear miniCic was quantified similarly from a single z slice.

Quantification of photoconversion experiments

Images were maximum projected over 2-3 z slices that contained the nucleus and then the background was subtracted using rolling ball subtraction with radius 50. A quantification box was drawn spanning the region 25-50 µm away from the illumination boundary. The level of red tdEOS fluorescence in that box was measured at each timepoint. Then the fold change in red fluorescence compared to the timepoint immediately following the illumination (0 minutes) was determined.

**Table S1: Primers used for cloning LlamaTag constructs**

| Primer ID | Sequence |
|---|---|
| TllN_left_Forward | gggcgaattgaatttagcggccgcgAAACGCAATCTGAGCTCCGC |
| TllN_left_Reverse | agccgccggaacctccagatccaccCATACCGATGTGAGGCGTAATTTTG |
| TllN_right_Forward | AggttctggtggttcaggaggttccCAGTCGTCGGAGGGTTCACC |
| TllN_right_Reverse | ctagtcctgcaggtttaaacgaattAGCTGGGCGAAATTCATGGG |
| HkbN_left_Forward | gggcgaattgaatttagcggccgcgAAGTTTAGGCGAACTGTACC |
| HkbN_left_Reverse | agccgccggaacctccagatccaccCATTTTGCAGTAGTTTTAATGGAAG |
| HkbN_right_Forward | AggttctggtggttcaggaggttccTCGACGATTAAGCTGCATCC |
| HkbN_right_Reverse | ctagtcctgcaggtttaaacgaattTGTTTAAAGCGTGCAAGCGC |
| Hkb_Q5_Forward | CTCGACGATTaatctgCATCCCCCGC |
| Hkb_Q5_Reverse | GAACCTCCTGAACCACCAGAACC |
| TllN_gRNA_Forward | GGACGACTGCATACCGATGTGTTTTAGAGCTAGAAATAGCAAG |
| TllN_gRNA_Reverse | GAAGTATTGAGGAAAACATACCTATATAAATG |
| HkbN_gRNA_Forward | GGTAGGTTTGCGGGGGATGCGTTGTTTTAGAGCTAGAAATAGCAAG |
| HkbN_gRNA_Reverse | GAAGTATTGAGGAAAACATACCTATATAAATG |

#### Legends for Supplemental Movies

**Movie S1: NC14 *byn*-MS2 dynamics under high and low light**
Movie shows *byn*-MS2 bursts during NC14 under high (top) and low (bottom) blue light. On the left are the raw images and on the right are segmented images. Orange nuclear masks show nuclei with a burst in corresponding frames. Timer (mm:ss) shows time since the start of NC14. Scale bar is 10 μm.

**Movie S2: *tll*- and *hkb*-MS2 dynamics under high and low light**
Sequential movies show *tll*-MS2 under high light, *tll*-MS2 under low light, *hkb*-MS2 under high light, and *hkb*-MS2 under low light. Each frame is labeled with the corresponding nuclear cycle (NC10-14). Scale bar is 10 μm. Timer (hh:mm:ss) shows time since the blue light was applied.

**Movie S3: NC14 *byn*-MS2 dynamics under high, short light**
Movie shows *byn*-MS2 bursts during NC14 when high light is applied only from NC10-NC13. Orange nuclear masks show nuclei with a burst in corresponding frames. Timer (mm:ss) shows time since the start of NC14. Scale bar is 10 μm.

**Movie S4: NC14 *tll*- and *hkb*-MS2 dynamics when light is applied at different developmental times**
Sequential movies show *tll*-MS2 bursts in NC14 and then *hkb*-MS2 bursts in NC14 when high light is applied starting in either NC13 (top) or NC14 (bottom). Teal and magenta nuclear masks show nuclei with a burst in corresponding frames. Timer (mm:ss) shows time since the start of NC14. Scale bar is 10 μm.

**Movie S5: NC14 *byn*-MS2 dynamics under high in illuminated and unilluminated regions**
Movie shows *byn*-MS2 bursts during NC14 when high light is applied to the illuminated region indicated by the blue bar. The dashed line indicates the illumination boundary. Orange nuclear masks show nuclei with a burst in corresponding frames. Timer (mm:ss) shows time since the start of NC14. Scale bar is 10 μm.
